# Testing the predictability of morphological evolution in contrasting thermal environments

**DOI:** 10.1101/609933

**Authors:** Natalie Pilakouta, Joseph L. Humble, Iain D.C. Hill, Jessica Arthur, Ana P.B. Costa, Bethany A. Smith, Bjarni K. Kristjánsson, Skúli Skúlason, Shaun S. Killen, Jan Lindström, Neil B. Metcalfe, Kevin J. Parsons

**Affiliations:** Institute of Biodiversity, Animal Health, and Comparative Medicine, University of Glasgow, Glasgow, UK; School of Biological Sciences, University of Aberdeen, Aberdeen, UK; School of Life Sciences, University of Nottingham, Nottingham, UK; Department of Aquaculture and Fish Biology, Hólar University, Sauðárkrókur, Iceland; Icelandic Museum of Natural History, Reykjavík, Iceland

**Author notes:** Corresponding authors: Natalie Pilakouta; Kevin Parsons.

**Keywords:** body shape, climate change, gene flow, geometric morphometrics, *Gasterosteus aculeatus*, parallel evolution, sympatric divergence, temperature, threespine stickleback

## Abstract

In light of climate change, the ability to predict evolutionary responses to temperature changes is of central importance for conservation efforts. Prior work has focused on exposing model organisms to different temperatures for just one or a few generations under laboratory conditions. Using a ‘natural experiment’, we show that studying parallel evolution in wild populations from contrasting thermal environments presents a more powerful approach for understanding and predicting responses to climate change. More specifically, we used a unique study system in Iceland, where freshwater populations of threespine sticklebacks (*Gasterosteus aculeatus*) are found in waters warmed by geothermal activity, adjacent to populations in ambient-temperature water. We used three sympatric and three allopatric warm-cold population pairs to test for repeated patterns of morphological divergence in relation to thermal habitat. We found that thermal habitat explained over 50% of body shape variation: fish from warm habitats had a deeper mid-body, a shorter jaw, and smaller eyes. Our common garden experiment showed that most of these morphological differences between thermal habitats were heritable. Lastly, absence of gene flow seems to facilitate parallel divergence across thermal habitats: all three allopatric population pairs were on a common evolutionary trajectory, whereas sympatric pairs followed different trajectories. Our findings therefore suggest that morphological responses to rising temperatures can be predictable when there is limited gene flow. On the other hand, migration of individuals between different thermal habitats or microhabitats can exaggerate nonparallel evolution and reduce our ability to predict evolutionary responses.

## INTRODUCTION

Understanding whether populations evolve in a predictable manner when exposed to similar environmental conditions is crucial for understanding adaptation. Studies on a wide range of taxa (insects: Nosil et al. 2002; fishes: Bernatchez et al. 2010; birds: Mundy 2005; mammals: Hoekstra 2006) have shown that different populations in similar environments tend to evolve similar phenotypes (Losos 2011). This pattern is referred to as parallel evolution and can arise due to natural selection and developmental bias. These evolutionary forces can result in evolutionary change that is predictable and repeatable (Schluter and Nagel 1995, Brakefield 2006, Oke et al. 2017, Uller et al. 2018).

Studying parallel evolution is especially valuable in light of global environmental change (de Amorim et al. 2017), because the ability to predict general population responses to anthropogenic change is of central importance for management and conservation efforts. In the coming decades, climate change will arguably pose the most significant threat to biodiversity. Rising temperatures are already altering abiotic and biotic environmental conditions and imposing novel selection pressures on organisms (Crozier & Hutchings 2014). Ectotherms, such as fishes and reptiles, are particularly vulnerable because of their high sensitivity to temperature changes (Zuo et al. 2012). Consequently, there is now a pressing need to understand the scope for populations to respond to climate change.

Although studies on plastic (within-generation) responses to temperature are rapidly accumulating, studies on the long-term evolutionary responses to climate change are still lacking (Crozier & Hutchings 2014). The few studies examining evolutionary responses have used laboratory experiments where model organisms were exposed to different temperature treatments over several generations (e.g., Alton et al. 2017). This approach can only examine the direct effects of temperature, but under natural conditions, changes in temperature will be accompanied by changes in various ecological factors, such as food availability, parasitism, and predation pressure (Crozier & Hutchings 2014). We therefore propose that studying parallel evolution in natural populations inhabiting contrasting thermal environments provides a more powerful approach for understanding and predicting population responses to increasing temperatures.

To this end, we took advantage of a ‘natural experiment’ in Iceland, where freshwater populations of threespine sticklebacks (*Gasterosteus aculeatus*) are found in waters warmed by geothermal activity (warm habitats), adjacent to populations in ambient-temperature water (cold habitats). This unique study system provides repeated and independent examples of populations inhabiting long-term contrasting thermal environments over a small geographic scale, thereby avoiding the confounding factors associated with latitudinal or elevational comparisons. Another notable attribute of this study system is that while most of these warm and cold habitats are in separate water bodies (allopatric), some are found in different parts of the same water body (sympatric). Movement of individuals, and thus gene flow, is possible between sympatric but not allopatric populations. This allows us to examine whether potential gene flow might influence the magnitude and/or direction of divergence between warm and cold habitats (hereafter referred to as thermal divergence). Lastly, the age of these warm habitats, and hence the maximum time these populations have experienced elevated temperatures, ranges from decades to thousands of years (Table 1). These different timescales make it possible to examine whether populations exposed to a warm environment for a relatively short time have diverged to the same extent as much older populations.

**Table 1.** Sampling locations and sample sizes of warm- and cold-habitat sticklebacks collected in May−June 2016. All cold habitats have existed since the last glacial period and are therefore approximately 10,000 years old, whereas warm habitats can be classified as either young (<100 years old) or old (>1,000 years old). Distance refers to how far apart the warm-habitat and cold-habitat sampling sites are for each population pair. The water temperature listed is the average temperature recorded at each sampling location during the summer.

| Population pair | Connectivity | Age of warm habitat | Distance (km) | Water body | Thermal habitat | Water temperature (°C) | Number of wild-caught specimens | Number of F1 specimens | Number of F1 families |
| --- | --- | --- | --- | --- | --- | --- | --- | --- | --- |
| A1 | Allopatric | Young | 0.03 | Unnamed | Warm | 22.4 | 29 | 40 | 8 |
|  |  |  |  | Unnamed | Cold | 14.0 | 31 | 40 | 9 |
| A2 | Allopatric | Old | 21.04 | Grettislaug | Warm | 24.9 | 29 | 40 | 5 |
|  |  |  |  | Garðsvatn | Cold | 14.6 | 30 | 40 | 8 |
| A3 | Allopatric | Old | 6.20 | Unnamed | Warm | 27.0 | 14 |  |  |
|  |  |  |  | Unnamed | Cold | 13.0 | 18 |  |  |
| S1 | Sympatric | Old | 3.18 | Mývatn | Warm | 22.8 | 30 | 40 | 7 |
|  |  |  |  |  | Cold | 11.5 | 30 | 40 | 7 |
| S2 | Sympatric | Young | 0.03 | Áshildarholtsvatn | Warm | 24.1 | 30 | 40 | 5 |
|  |  |  |  |  | Cold | 12.2 | 30 | 40 | 8 |
| S3 | Sympatric | Young | 0.10 | Húseyjarkvísl | Warm | 23.9 | 28 |  |  |
|  |  |  |  |  | Cold | 10.3 | 32 |  |  |

Here, we focus on temperature-driven evolution in morphology. Morphology can determine fitness by influencing reproduction, foraging ability, and swimming performance (Rowinski et al. 2015). Morphological variation is also related to sexual selection and reproductive isolation and can therefore contribute to population divergence and speciation (Head et al. 2013). Previous research has shown that morphology often exhibits similar patterns of adaptive divergence across populations in response to common environmental conditions (Jastrebski and Robinson 2004, Cooper et al. 2010). Rearing temperature, in particular, directly influences the development of body shape in fishes (Sfakianakis et al. 2011, Ramler et al. 2014, Rowinski et al. 2015). Yet, it is still unknown whether evolutionary responses to temperature changes are repeatable and thus predictable.

Our study addresses this gap in our knowledge by testing whether there is morphological divergence between sticklebacks from six warm-cold population pairs, and if so, whether it follows parallel patterns. Evidence for parallelism would suggest that responses to elevated temperature are predictable. Nevertheless, even if our six warm-cold population pairs share evolutionary trajectories, they may not necessarily show the same degree of divergence. Hence, we also investigated whether the magnitude of thermal divergence differs across population pairs, as a function of population age or habitat connectivity (i.e., potential for gene flow). Lastly, to determine whether morphological differences observed in wild-caught sticklebacks are heritable, we conducted a common garden experiment by breeding fish from warm and cold habitats and rearing their offspring at a common temperature. By addressing these questions, we can advance our ability to predict evolutionary responses to elevated temperatures in light of global climate change.

## METHODS

### Collecting wild-caught sticklebacks

We used unbaited minnow traps to collect adult threespine sticklebacks from six warm-cold population pairs in Iceland in May−June 2016 (Table 1, Figure 1). Three of these population pairs were allopatric (designated A1-3), meaning that the warm and cold habitats were in neighbouring but separate water bodies with no current potential for gene flow (Table 1). The other three population pairs were sympatric (designated S1-3), meaning that the warm and cold habitats were found in the same water body with no physical barriers between them and thus a potential for gene flow (Table 1). The cold habitats have all existed since the last glacial period about 10,000 years ago (Einarsson et al. 2004), but there is some variation in the age of the warm habitats (Table 1). The A1, S2, and S3 warm habitats originated 50−70 years ago and are fed by excess hot water runoff from nearby residences that use geothermal heating. The remaining warm habitats have been naturally heated by geothermal vents for over 1,000 years (Hight 1965, Einarsson et al. 2004).

**Figure 1.**
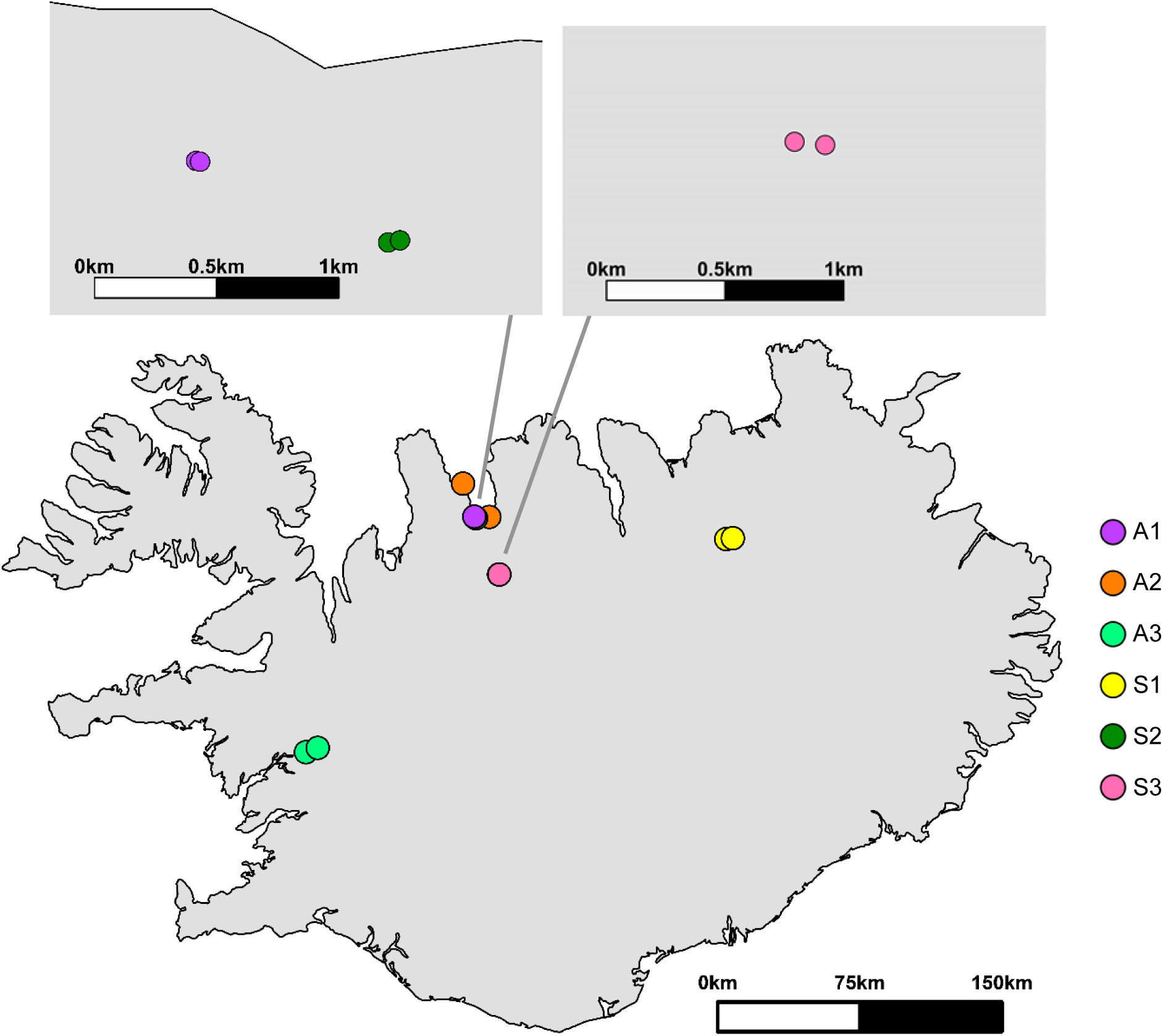
Map of Iceland showing the sampling locations of warm- and cold-habitat sticklebacks we collected for this study. All sticklebacks were collected from freshwater populations, and each of the six population pairs (A1, A2, A3, S1, S2, and S3) is indicated by a different colour.

A subset of sticklebacks (*n*=331) caught in minnow traps were immediately euthanised using an overdose of phenoxyethanol and preserved in 10% buffered formalin (Table 1). In addition, approximately 100 sticklebacks from each of eight sampling locations (i.e., four warm-cold population pairs) were kept in temporary holding tanks at Hólar University before being transported to University of Glasgow to be used for breeding in our common garden experiment (see below).

### Transport of study animals and animal husbandry

We fasted sticklebacks for 48 hr to minimise the build-up of ammonia in the transport water. On the day of shipping, we placed approximately 100 sticklebacks from each population in 100-litre polyethylene bags containing 25 litres of water. Air was removed from the bags and replaced with pure oxygen. Bags were sealed and placed inside insulated Styrofoam shipping boxes to minimise temperature fluctuations during transport. The fish were in transit for approximately 72 hr before arriving in Glasgow. No mortality was observed during transport.

Once the fish arrived at the University of Glasgow, they were kept at densities of 10-15 individuals per 10-litre tank in a common recirculation system at 15°C. This intermediate temperature is close to the maximum temperature experienced by fish in cold habitats in the summer and the minimum temperature experienced by fish in warm habitats in the winter (Pilakouta et al. 2020). All tanks contained plastic plants as shelter and air stones to oxygenate the water. Fish were fed *ad libitum* twice a day with a mixture of frozen bloodworms, *Mysis* shrimp, and *Daphnia*. They were kept at a 12 hr light:12 hr dark photoperiod.

### Common garden experiment

We carried out a common garden experiment to determine whether morphological variation between warm and cold habitats was heritable. For this experiment, we bred wild-caught sticklebacks from warm and cold habitats of two allopatric population pairs (A1 and A2) and two sympatric population pairs (S1 and S2). Gravid females and males displaying breeding colours were euthanised with an overdose of benzocaine and used for *in vitro* fertilisation (Barber & Arnott 2000). After performing *in vitro* fertilisation in petri dishes, we placed fertilised embryos in mesh baskets submerged in well-aerated water with methylene blue (2.5 μg/ml) until hatching. These F1 generation stickleback larvae were fed with newly hatched HUFA-enriched *Artemia salina* nauplii, microworms, and powdered food (ZM100 and ZM200 fry food, ZM Systems, Twyford, UK) until large enough to eat pelleted food (Microstart, EWOS Ltd, Surrey, UK) at a standard length of approximately 2 cm. At that stage, they were transferred to 10-litre tanks and kept at densities of 15-20 individuals. They were maintained at a constant water temperature of 18°C (±0.5°C) from the embryonic stage to adulthood. About 12 months after hatching, we euthanised 320 of these F1 individuals using an overdose of benzocaine and preserved them in 10% buffered formalin (*n*=40 per sampling location from at least 5 full-sib families; Table 1).

### Specimen preparation

All preserved specimens were bleached and cleared to remove skin pigmentation and make the body translucent (Potthoff 1984). They were then stained with Alizarin Red S to emphasize bone morphology and were stored in 75% glycerol until excess stain was removed. Individual specimens (*n*=331 wild-caught sticklebacks, *n*=320 F1 sticklebacks) were photographed on their left side with a Canon EOS 1100D digital camera (Canon Inc, Tokyo, Japan). All photographs included a scale and were taken from a fixed distance and angle using a copy stand.

### Linear measurements

We used the photographs of wild-caught and F1 sticklebacks to measure pectoral fin length and dorsal spine length, which are related to swimming and defence, respectively (Drucker et al. 2005, Hoogland et al. 1956). Using the software program tpsDig2 (Rohlf 2018), we placed landmarks on the base and tip of the longest fin ray (pectoral fin length) and on the base and tip of the first and second spine (dorsal spine lengths). We then calculated interlandmark distances using CoordGen8 (Zelditch et al. 2012) and regressed each length measurement against centroid size to minimise body size and allometric effects. Because they belong to articulated structures, these landmarks were only used for obtaining linear distances and were not included in the multivariate body shape analysis.

### Body shape variation

To measure body shape in wild-caught and F1 sticklebacks, we placed 22 anatomical landmarks and 10 sliding semilandmarks (Bookstein 1997) on each image to quantify variation in the lateral view using a geometric morphometric approach (Figure 2). Sliding semilandmarks are points along curves that measure variation that cannot be captured by anatomical landmarks. We placed 10 equally spaced sliding landmarks between the anterior tip of the upper jaw and the posterior tip of the frontal bone to measure variation in head curvature (Figure 2). The sliding procedure was conducted based on chord distance using the tpsRelw32 software (Rohlf 2019).

**Figure 2.**
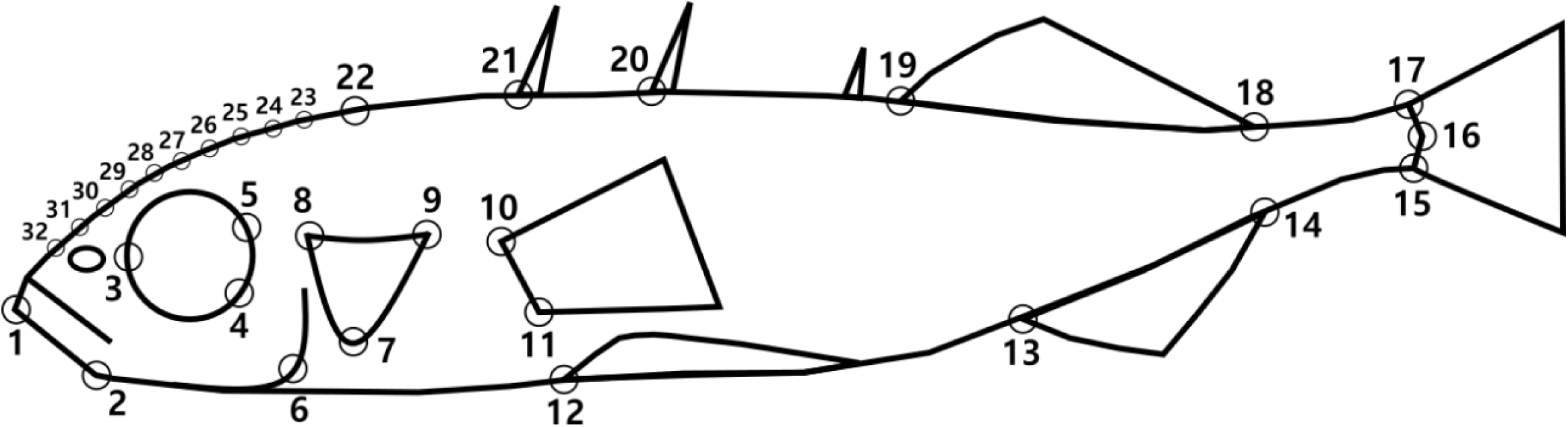
Depiction of the anatomical landmarks (large open circles) and sliding landmarks (small open circles) used to compare the body shape of sticklebacks from cold and warm habitats. The anatomical landmarks were placed on the lower jaw (1, 2), eye orbit (3−5), preopercle (6), opercle (7−9), pectoral fin insertion points (10, 11), anterior tip of pelvic spine (12), anal fin insertion points (13, 14), caudal fin insertion points (15, 17), caudal border of hypural plate at lateral midline (16), dorsal fin insertion points (18, 19), dorsal spine insertion points (20, 21), and the posterior tip of the frontal bone (22). The sliding landmarks were placed between the tip of the upper jaw and the posterior tip of the frontal bone (23−32) to examine head curvature.

To reduce the effects of size and orientation across individuals, we performed a Generalized Procrustes Analysis using CoordGen8 (Rohlf and Slice 1990). This process superimposes landmark configurations to minimise the sum of squared distances between corresponding landmark configurations by scaling, rotating, and translating specimens in relation to their geometric centre. To minimise the potential effects of allometry on the data, we used Standard6 (Zelditch et al. 2012) to perform a multiple regression of shape on geometric centroid size to generate residuals. We then performed a thin-plate spline (TPS) procedure to generate partial warp scores for further statistical analysis. This procedure models the form of an infinitely thin metal plate that is constrained at some combination of points (i.e., landmarks) but is otherwise free to adopt a target form in a way that minimizes bending energy. In morphometrics, this interpolation is applied to a Cartesian coordinate system in which deformation grids are constructed from two landmark configurations (Bookstein 1991). The total deformation of the thin-plate spline (including uniform components) can be decomposed into geometrically orthogonal components (partial warps) based on scale (Rohlf and Marcus 1993). TPS and the generation of partial warps scores was performed using PCAgen (Zelditch et al. 2012).

### Data analysis

Analyses were run using R version 3.5.1 (R Core Team 2018) and figures were generated using the ggplot2 package (Wickham 2009) unless otherwise noted.

#### Linear measurements

For each wild-caught and F1 stickleback, we obtained linear distances for three morphological traits: pectoral fin length, first dorsal spine length, and second dorsal spine length. After checking for homogeneity of slopes, we regressed each linear distance against centroid size and used the residuals from these regressions in separate ANOVA models for each trait. The explanatory variables in these models were population pair, thermal habitat, and the interaction between population pair and thermal habitat. The main effect of population pair summarises properties unique to different replicates (Bolnick et al. 2018). The main effect of thermal habitat measures the extent to which thermal divergence is shared across replicate locations and thus measures parallel evolution (Bolnick et al. 2018). The population pair × thermal habitat interaction indicates how the direction and magnitude of thermal divergence varies among population pairs, implying nonparallel evolution. To determine the partial variance explained by each factor and interaction, we used the heplots package to calculate partial eta squared (η^2^) values (Fox et al. 2007).

#### Body shape variation

To test whether thermal habitat affects body shape, we performed a discriminant function analysis (DFA) for each population pair using thermal habitat (warm vs cold) as a grouping variable to explain variation in partial warp scores (i.e., body shape). This analysis allowed us to characterize the potentially divergent effects of temperature within each population pair. We then performed another DFA which included all population pairs together to examine the overall effect of temperature on body shape.

Next, we used a MANOVA model that included partial warp scores from all populations as the response variable. This model allowed us to separate the independent and interactive effects of thermal habitat (warm vs cold) and population pair on body shape. As mentioned above, a significant effect of thermal habitat would imply parallel evolution, whereas a significant effect of the thermal habitat × population pair interaction would imply nonparallel evolution. We again calculated partial eta squared (η^2^) values to determine the partial variance explained by each factor or interaction in this MANOVA model. Our MANOVA and DFA approaches modelled variation in body shape but did not allow us to separate the effects of direction and magnitude of thermal divergence, so we adopted two additional approaches.

#### Direction of thermal divergence in body shape

*—*To examine parallel evolution in wild-caught fish, we compared the scale-free vector of divergence for each population pair using the canonical scores derived from the DFA. To derive the vector of divergence, we regressed the Procrustes-superimposed landmark data from each population pair on its corresponding canonical axis. The observed angle between vectors for all pairwise comparisons of populations was then calculated as the arccosine. We ran 900 bootstraps with replacement for each population pair independently and calculated 95th percentiles of the range of angles obtained by resampling. To carry out the bootstrapping procedure, the two thermal groups were merged into a common pool, and two groups with the same sample size as the original data sets were drawn with replacement from the common pool.

The observed angle between two population pairs was compared against the angle within each pair to determine whether it differed from random processes (Zelditch et al. 2012). If the between-population-pair angle exceeded both of the within-population-pair angles, this meant that the population pairs were evolving along different evolutionary trajectories (Parsons et al. 2011, Zelditch et al. 2012, Parsons et al. 2016). These procedures were performed using the tool VecCompare in the software Regress8 (Zelditch et al. 2012).

#### Magnitude of thermal divergence in body shape

*—*To assess the magnitude of shape divergence between wild-caught fish from warm and cold habitats, we used a Procrustes distance-based approach, which allowed us to compare the six population pairs in a common shape space (i.e., a common scale). To this end, we calculated the Procrustes mean for the warm group and cold group in each population pair. Following the determination of observed distances in Procrustes means based on an *F*-value, we performed 900 bootstraps to determine the probability for it to have been produced by chance. As before, we carried out the bootstrapping procedure by merging the two thermal groups into a common pool; two groups with the same sample size as the original data sets were then drawn with replacement from the common pool. This analysis was carried out using the IMP TwoGroup8 software (Zelditch et al. 2012).

## RESULTS

### Linear measurements in wild-caught fish

Wild-caught sticklebacks from warm habitats had longer second (but not first) dorsal spines in most population pairs, as indicated by a statistically significant interaction between thermal habitat and population pair (Supplementary Table 1, Supplementary Figure 1). The effect of thermal habitat on pectoral fin length also varied across population pairs (Supplementary Table 1, Supplementary Figure 1). Sticklebacks from warm habitats had longer pectoral fins in some pairs (A2, S1) but shorter pectoral fins in other pairs (A1, S3).

### Body shape variation in wild-caught fish

The discriminant function analysis showed strong groupings based on thermal habitat in all six of our population pairs (97% correct classification for the A3 population pair and 100% correct classification for all other pairs), indicating that wild-caught fish could be reliably assigned to warm or cold habitats on the basis of their body shape (Figure 3). When all six population pairs were included together in the DFA, there was 82% correct classification based on thermal habitat (Figure 4). Overall, sticklebacks from warm habitats tended to have smaller eyes, a shorter jaw, and a deeper mid-body tapering to a narrower caudal peduncle (Figure 4).

**Figure 3.**
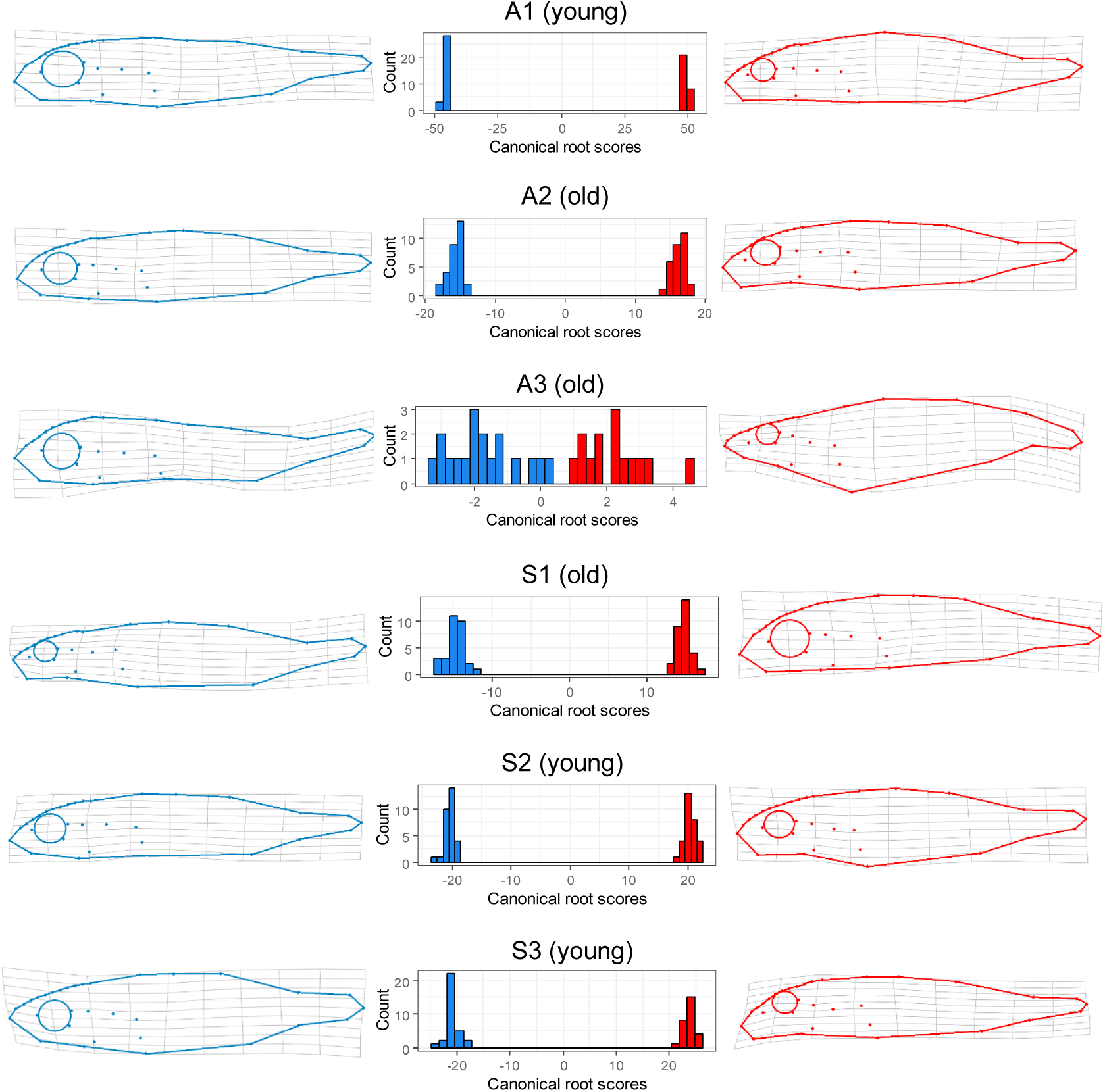
Thermal divergence in morphology of wild-caught sticklebacks from three allopatric population pairs (A1–A3) and three sympatric population pairs (S1–S3). Plots show frequency histograms of linear discriminant (LD1) scores from the DFA run on partial warp scores, along with thin plate spline deformations showing the observed extremes in each population pair. Wild-caught specimens from cold and warm habitats are indicated in blue and red, respectively. “Young” or “old” refers to the age of the warm habitat in that population pair. The deformation grids were generated using tpsRegr (Rohlf 2008). The shape differences were extrapolated by a factor of 3 to allow easier interpretation.

**Figure 4.**
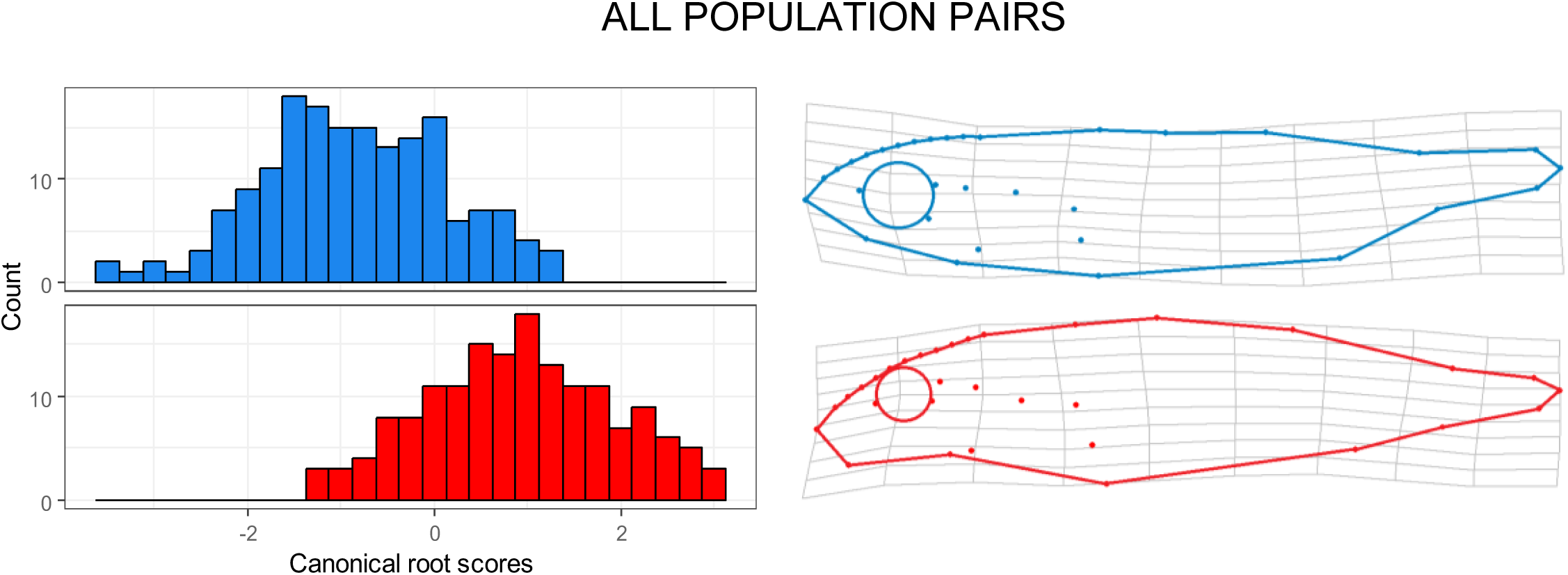
Thermal divergence in morphology of wild-caught sticklebacks pooled across all population pairs (A1, A2, A3, S1, S2, and S3). The left panel shows frequency histograms of linear discriminant (LD1) scores from the DFA run on partial warp scores, and the right panel shows thin plate spline deformations of the observed extremes. Specimens from cold and warm habitats are indicated in blue and red, respectively. The deformation grids were generated using tpsRegr (Rohlf 2008). The shape differences were extrapolated by a factor of 3 to allow easier interpretation.

The MANOVA model provided confirmation that differences in temperature have led to a divergence in body shape: thermal habitat had a significant effect on body shape across population pairs, explaining 53% of the observed variation in shape (Table 2). Body shape was also influenced by the interaction between thermal habitat and population pair, indicating that changes in body shape due to temperature varied across population pairs (Table 2).

**Table 2.** Results of MANOVA model testing the effects of thermal habitat, population pair, and their interaction on body shape (i.e., partial warp scores) of wild-caught sticklebacks and lab-reared F1 sticklebacks. Df denotes degrees of freedom. Statistically significant *P*-values are indicated in bold.

|  | Wild-caught sticklebacks |  |  |  |  | F1 generation sticklebacks |  |  |  |  |
| --- | --- | --- | --- | --- | --- | --- | --- | --- | --- | --- |
| | Df | Wilk's $\lambda$ | *F* | *P* | Partial variance explained ($\eta^2$) | Df | Wilk's $\lambda$ | *F* | *P* | Partial variance explained ($\eta^2$) |
| Thermal habitat | 1 | 0.471 | 4.96 | **<0.0001** | 53% | 1 | 0.320 | 8.97 | **<0.0001** | 68% |
| Population pair | 5 | 0.012 | 6.28 | **<0.0001** | 56% | 3 | 0.057 | 6.76 | **<0.0001** | 60% |
| Thermal habitat<br>× population pair | 5 | 0.033 | 4.38 | **<0.0001** | 47% | 3 | 0.077 | 5.69 | **<0.0001** | 55% |
| Error | 319 |  |  |  |  | 312 |  |  |  |  |

### Direction of thermal divergence in body shape of wild-caught fish

We found mixed evidence for parallelism in the divergence trajectories of our warm-cold population pairs (Table 3). The trajectories of thermal divergence were parallel for all comparisons between allopatric population pairs (A1−A2, A1−A3, and A2−A3) but for none of the comparisons between sympatric population pairs (S1−S2, S1−S3, S2−S3). For example, in all three allopatric population pairs, warm-habitat fish had smaller eyes and shorter jaws than cold-habitat fish (Figure 3). In contrast, in sympatric population pairs, eye size and jaw length did not seem to vary consistently based on thermal habitat (Figure 3).

**Table 3.** Vector analysis for body shape of wild-caught sticklebacks. We present data on the angle between each set of population pairs, as well as the 95th percentile of the range of angles (obtained by resampling) within each population pair. If the angle between two population pairs exceeds both of the angles within each population pair, we can conclude that those population pairs are evolving along different evolutionary trajectories. The results in bold indicate evidence for parallelism in thermal divergence between the two population pairs.

| Population pairs | Angle between population pairs | Angle within population pair 1 | Angle within population pair 2 |
| --- | --- | --- | --- |
| A1–A2 | **67°** | **42°** | **69°** |
| A1–A3 | **62°** | **84°** | **59°** |
| A2–A3 | **81°** | **101°** | **60°** |
| A1–S1 | 127° | 41° | 32° |
| A1–S2 | 93° | 42° | 64° |
| A1–S3 | 72° | 35° | 41° |
| A2–S1 | 112° | 70° | 31° |
| A2–S2 | 110° | 69° | 65° |
| A2–S3 | **50°** | **36°** | **70°** |
| A3–S2 | **73°** | **58°** | **94°** |
| A3–S3 | **81°** | **115°** | **59°** |
| A3–S1 | 122° | 60° | 61° |
| S1–S2 | 92° | 31° | 64° |
| S1–S3 | 116° | 35° | 32° |
| S2–S3 | 117° | 35° | 63° |

### Magnitude of thermal divergence in body shape of wild-caught fish

Consistent with the DFA and MANOVA results, the distance-between-means test provided strong evidence for divergence in body shape between sticklebacks from warm and cold habitats (Supplementary Table 1). The magnitude of thermal divergence varied across population pairs but did not seem to be related to population age or connectivity between warm and cold habitats (Supplementary Table 1).

### Linear measurements in lab-reared F1 fish

Lab-reared F1 sticklebacks from warm habitats had longer first dorsal spines in most population pairs, as indicated by a statistically significant effect of thermal habitat (Supplementary Table 3, Supplementary Figure 2). This demonstrated heritable differences in first dorsal spine length, but there was no such evidence for divergence in second dorsal spine length or pectoral fin length between thermal habitats.

### Body shape variation in lab-reared F1 fish

Lab-reared F1 sticklebacks from different thermal origins differed in body shape after being reared at the same temperature, demonstrating that heritable variation underlies their morphological divergence. Our discriminant function analysis showed strong groupings based on the parents’ thermal habitat in all four population pairs (99% correct classification for the S1 population pair and 100% correct classification for the remaining pairs). These results indicate that lab-reared F1 fish could be reliably assigned to their parents’ thermal habitat of origin on the basis of their body shape (Figure 5; Supplementary Figure 3). When all four population pairs were included together in the DFA, there was 93% correct classification based on their parents’ thermal habitat of origin.

**Figure 5.**
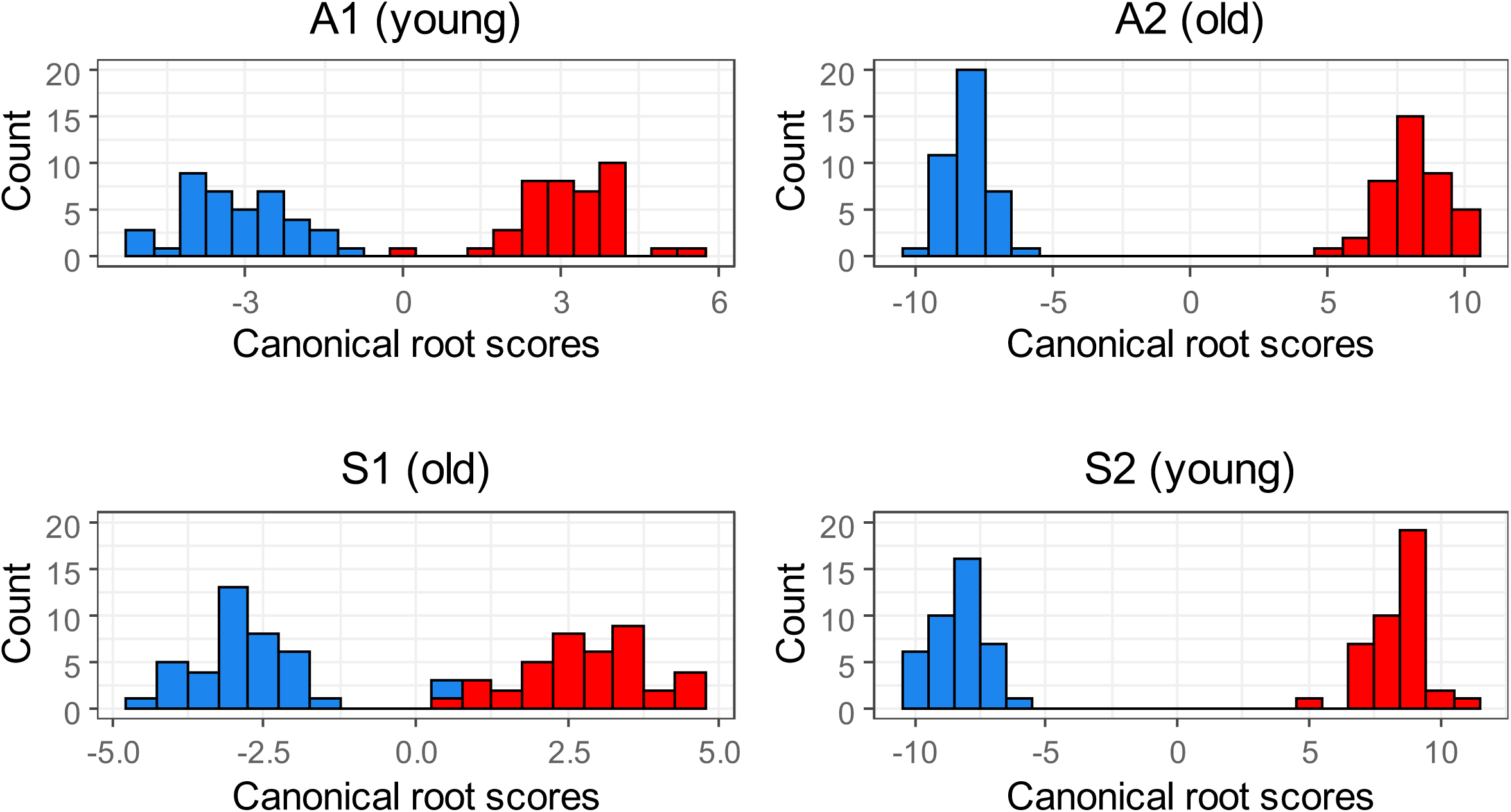
Thermal divergence in morphology of lab-reared F1 sticklebacks from four population pairs (A1, A2, S1, and S2). Plots show frequency histograms of linear discriminant (LD1) scores from the DFA run on partial warp scores. These F1 generation sticklebacks were all reared at a common temperature (18°C). Blue and red bars represent specimens whose parents were collected from cold and warm habitats, respectively. “Young” or “old” refers to the age of the warm habitat in that population pair.

The MANOVA model provided confirmation for shared evolutionary differences in morphology: thermal habitat of origin had a significant effect on body shape across population pairs, explaining 68% of the observed variation in shape (Table 2). Body shape was also influenced by the interaction between thermal habitat and population pair, reflecting variation in thermal divergence among population pairs (Table 2). Most morphological differences observed in the wild-caught fish persisted under a common rearing temperature, with warm-origin F1 fish having a deeper mid-body, a deeper caudal peduncle, and a steeper craniofacial profile than cold-origin F1 fish.

## DISCUSSION

Here, we tested for morphological divergence between sticklebacks from different thermal habitats and examined whether such divergence follows parallel patterns across population pairs. We found evidence for shared differences in body shape between wild-caught sticklebacks from warm and cold habitats, with temperature explaining over half of this variation. Our common garden experiment showed that these morphological differences are strongly influenced by heritable variation and thus reflect a common evolutionary response to the contrasting thermal environments these fish inhabit. Population age did not seem to influence the magnitude or direction of thermal divergence in wild-caught sticklebacks, but connectivity between thermal habitats did seem to influence the direction of the divergence: allopatric population pairs were on parallel evolutionary trajectories, whereas sympatric pairs were on different trajectories.

### Effects of thermal environment on morphological evolution

It is well established that water temperature can influence body shape development in fishes through plastic responses (Marcil et al. 2006, Sfakianakis et al. 2011, Ramler et al. 2014, Rowinski et al. 2015). Generally, higher temperatures lead to increased body depth (Marcil et al. 2006, Sfakianakis et al. 2011, Rowinski et al. 2015). Consistent with this, we found that in most population pairs, wild-caught sticklebacks from warm habitats were more deep-bodied with a narrower caudal peduncle than those from cold habitats. They also tended to have smaller eyes, a shorter lower jaw, and a longer second dorsal spine (Figure 3).

Although our study presents strong evidence for heritable morphological differences between sticklebacks from different thermal habitats, it is unclear whether these differences are adaptive, and if so, what the underlying causes are. Since the effects of temperature on body shape can be either direct or indirect (i.e., mediated by changes in other ecological conditions), the observed morphological divergence between thermal habitats could be due to selection from such indirect effects. For example, changes in jaw length and body depth may be driven by differences in food availability (Rowinski et al. 2015) or diet composition (Hjelm et al. 2001). More specifically, greater body depth can be related to jaw function, as it signifies hypertrophied epaxial musculature. This suggests an increased suction ability and, along with a shorter jaw, would indicate a more benthic foraging ecology in the warm-habitat sticklebacks (McGee et al. 2013).

Another potential explanation for our findings is that sticklebacks in warm habitats have evolved deeper bodies in response to a higher predation risk. Greater body depth is thought to improve predator escape performance through increased manoeuvrability or predator gape limitation (Reimchen 1991, Walker 1997, Dominici et al. 2008). Similarly, dorsal spines are an antipredator defence and are generally longer in populations that experience elevated predation pressure (e.g., Blouw and Hagen 1984). Here, we found evidence for longer second dorsal spines in wild fish from warm habitats but longer first dorsal spines in lab-reared F1 fish from warm habitats. This suggests that evolved differences in the developmental system between thermal habitats keep the first spine the same under natural conditions but allow the second dorsal spine to respond via plasticity. In our study system, we expect bird predation to be higher in warm habitats, due to the lack of ice cover during the winter and the fact that birds tend to be attracted to warmer areas (Rowiniski et al. 2015). On the other hand, sticklebacks in warm habitats likely experience a lower risk of predation from freshwater piscivorous fish, which may be unable to cope with high temperatures (Eliason et al. 2011). Further research will be needed to investigate the functional significance of the morphological differences we have documented.

### Magnitude of thermal divergence in body shape

The magnitude of divergence between populations in warm and cold habitats may be influenced by various factors, such as connectivity between these habitats. For example, some researchers have argued that gene flow will constrain divergence between ecotypes when there is a potential for physical dispersal (Slatkin 1985, Lenormand 2002, Hendry and Taylor 2004). Under this scenario, we would expect sympatric population pairs to be less divergent than allopatric population pairs (Hendry and Taylor 2004, Pinho and Hey 2010). However, sympatric pairs could instead be more divergent because of character displacement, whereby differences between ecotypes are more pronounced in areas where they co-occur and minimised in areas where their distributions do not overlap. This pattern results from trait evolution driven by competition among ecotypes, or closely related species, for a limited resource (Brown and Wilson 1956, Losos 2011).

In our study, the presence or absence of geographical barriers did not seem to influence the magnitude of thermal divergence in body shape. It is interesting to note that despite the potential for gene flow in sympatric population pairs, we found no evidence for intermediate phenotypes (Figure 3). This was particularly surprising for sympatric pairs S2 and S3, where the warm and cold habitats are only 30 and 100 meters apart, respectively. In fact, the only population pair where we observed some overlap in the phenotypic distribution of body shape was an allopatric pair. There are several plausible explanations for the absence of intermediate phenotypes in sympatric population pairs, including strong performance trade-offs, assortative mating, and hybrid inviability (Maynard Smith 1966, Schluter 2009).

The magnitude of thermal divergence in morphology may also vary depending on population age. In populations that have been diverging for longer, there is more scope for natural selection and genetic drift to introduce adaptive or stochastic phenotypic differences (Ord and Summers 2015). In our study system, we would thus expect relatively young population pairs (<100 years old) to be less divergent than old population pairs (>1,000 years old). There was no such evidence based on the observed morphological differences in wild-caught fish: populations in warm habitats that have only existed for a few decades were equally divergent from the corresponding cold populations as populations in warm habitats that have existed for thousands of years. This suggests that elevated temperature may drive rapid morphological changes, which are then relatively stable over a prolonged evolutionary timescale (Stockwell et al. 2003).

Although there was no indication that the magnitude of thermal divergence was related to population age in wild-caught fish, there was some indication of a greater degree of differentiation in older populations in the lab-reared F1 fish (Supplementary Figure 3). Nevertheless, we are cautious to draw strong conclusions from this comparison given that we only have data for two old and two young population pairs in our common garden experiment. Data on additional population pairs, including their within-generation responses to temperature, would be useful for addressing the idea that young populations are converging on similar phenotypes to older populations through phenotypic plasticity.

### Direction of thermal divergence in body shape

Habitat connectivity may influence not only the magnitude of divergence but also the degree of parallelism between replicate populations (Bolnick et al. 2018). Gene flow between different habitat types is thought to constrain local adaptation within each habitat, so if there is variation in the extent of gene flow among replicate populations, migration-selection balance will act differently contributing to nonparallel evolution (Hendry and Taylor 2004, Moore et al. 2007, Stuart et al. 2017). As a result, allopatric population pairs (no gene flow) and sympatric population pairs (potential for gene flow) may differ in their degree of parallelism. Indeed, we found that the absence of gene flow facilitates parallel divergence in warm-cold population pairs of sticklebacks: allopatric population pairs have all evolved along parallel trajectories, whereas none of the sympatric population pairs share evolutionary trajectories.

Previous theoretical and empirical work suggests that the degree of parallelism between replicate populations may also depend on the duration of evolutionary divergence (Lucek et al. 2014, Ord and Summers 2015). For example, population pairs that have been diverging for longer have more scope for natural selection and genetic drift to alter their evolutionary trajectories, resulting in a lower degree of parallelism in older populations (Bolnick et al. 2018). Yet, if evolution is limited by mutation rate, older populations will have had more time to accumulate similar adaptive mutations that produce a similar phenotypic solution in response to a particular environment (Orr 2005, Whitlock and Gomulkiewicz 2005). Under this scenario, older populations would have a higher degree of parallelism. Our findings did not support either of these possible outcomes, since the extent of parallel divergence did not seem to differ between young and old population pairs.

Lastly, it should be noted that even though we focused on the effects of habitat connectivity and population age, several other factors can influence the magnitude and direction of divergence in wild populations. These include ancestry and evolutionary history (Langerhans and DeWitt 2004), initial and ongoing effective population sizes (Szendro et al. 2013), variation in sexual selection (Bonduriansky 2011, Maan and Seehausen 2011), and many-to-one mapping, which refers to multiple phenotypic solutions to the same functional problem (Gould and Lewontin 1979, Wainwright et al. 2005). Further work will be needed to examine these additional factors.

### Predictability of evolution and adaptation to climate change

As discussed above, all the allopatric population pairs in this study share a similar divergence trajectory in terms of their thermal divergence in body shape. This parallelism could be due to natural selection, developmental bias, or their interaction (Losos 2011, Brakefield 2006, Uller et al. 2018). We cannot separate the effects of these processes in the present study, but regardless of the underlying causes, our results suggest that the absence of gene flow facilitates parallel evolution between warm and cold populations.

Thus, morphological evolution in response to increasing temperatures could be predictable to some extent for fish populations where there is little to no gene flow from other thermal habitats. Under these conditions, we may expect fish to evolve shorter jaws and a deeper mid-body after being exposed to elevated temperatures over multiple generations. On the other hand, migration of individuals between different thermal habitats or microhabitats will exaggerate nonparallel evolution (Oke et al. 2017, Bolnick et al. 2018) and reduce our ability to predict evolutionary responses to changes in temperature.

## CONCLUSION

Studying parallel evolution in natural populations inhabiting contrasting thermal environments presents a powerful approach for understanding and predicting population responses to increasing temperatures. Here, we have taken advantage of a unique study system that provides repeated and independent examples of populations found in different thermal environments in the absence of latitudinal or elevational variation. We show that, in populations with no gene flow from other thermal habitats, it can be possible to predict morphological evolution in response to elevated temperatures. Our findings therefore provide novel insights into how gene flow might influence temperature-driven parallel evolution and how fish populations may adapt to a warming world.

## ACKNOWLEDGEMENTS

We would like to thank Tiffany Armstrong, Anna Persson, and Kári Heidar Árnason for their help with fieldwork in Iceland, and Peter Koene for his help with photographing specimens. We are also grateful to Graham Law, Alistair Kirk, and Ross Philips for their assistance with animal husbandry and aquarium maintenance.

## ETHICAL STATEMENT

Our study adheres to the ASAB/ABS Guidelines for the Use of Animals in Research, the institutional guidelines at University of Glasgow, and the legal requirements of the UK Home Office (Project License P89482164).

