## Supplementary for "Testing the predictability of morphological evolution in contrasting thermal environments"

**Supplementary Table 1.** Results of ANOVA models testing the effects of thermal habitat (warm vs cold), population pair, and the interaction between thermal habitat and population pair on first dorsal spine length, second dorsal spine length, and pectoral fin length in wild-caught sticklebacks. Df denotes degrees of freedom. Partial eta-squared ($\eta^2$) values provide an estimate for partial variance explained by each factor or interaction. Statistically significant *P*-values are indicated in bold.

| | First dorsal spine length | | | | Second dorsal spine length | | | | Pectoral fin length | | | |
|---|---|---|---|---|---|---|---|---|---|---|---|---|
| | Df | *F* | *P* | Partial variance explained ($\eta^2$) | Df | *F* | *P* | Partial variance explained ($\eta^2$) | Df | *F* | *P* | Partial variance explained ($\eta^2$) |
| Thermal habitat | 1 | 0.16 | 0.69 | <0.1% | 1 | 24.8 | **<0.001** | 7.0% | 1 | 0.06 | 0.80 | <0.1% |
| Population pair | 5 | 1.24 | 0.29 | 1.9% | 5 | 2.46 | **0.033** | 3.9% | 5 | 8.59 | **<0.001** | 12.4% |
| Thermal habitat × population pair | 5 | 1.06 | 0.38 | 1.6% | 5 | 9.65 | **<0.001** | 13.0% | 5 | 3.44 | **0.005** | 5.4% |
| Error | 319 | | | | 319 | | | | 319 | | | |

**Supplementary Table 2.** Distances-between-means test comparing the magnitude of divergence in body shape between wild-caught sticklebacks from warm vs cold habitats in each population pair. "Young" or "old" refers to the age of the warm habitat in that population pair. Statistically significant *P*-values are indicated in bold.

| Population pair | *F* | *P* | Distance between means |
|---|---|---|---|
| A1 (young) | 6.49 | **0.001** | 0.022 |
| A2 (old) | 1.90 | 0.082 | 0.011 |
| A3 (old) | 2.71 | **0.039** | 0.024 |
| S1 (old) | 12.57 | **0.001** | 0.032 |
| S2 (young) | 2.55 | **0.030** | 0.014 |
| S3 (young) | 9.48 | **0.001** | 0.028 |

**Supplementary Table 3.** Results of ANOVA models testing the effects of thermal habitat (warm vs cold), population pair, and the interaction between thermal habitat and population pair on first dorsal spine length, second dorsal spine length, and pectoral fin length in F1 sticklebacks reared under a common temperature of 18˚C. Df denotes degrees of freedom. Partial eta-squared ($\eta^2$) values provide an estimate for partial variance explained by each factor or interaction. Statistically significant *P*-values are indicated in bold.

| | First dorsal spine length | | | | Second dorsal spine length | | | | Pectoral fin length | | | |
|---|---|---|---|---|---|---|---|---|---|---|---|---|
| | Df | *F* | *P* | Partial variance explained ($\eta^2$) | Df | *F* | *P* | Partial variance explained ($\eta^2$) | Df | *F* | *P* | Partial variance explained ($\eta^2$) |
| Thermal habitat | 1 | 9.05 | **<0.001** | 2.8% | 1 | 1.90 | 0.169 | 0.6% | 1 | 0.01 | 0.920 | <0.1% |
| Population pair | 1 | 0.09 | 0.770 | <0.1% | 1 | 1.07 | 0.302 | 0.3% | 1 | 0.526 | 0.469 | 0.2% |
| Thermal habitat x population pair | 1 | 1.21 | 0.271 | 0.4% | 1 | 0.02 | 0.904 | | 1 | 0.874 | 0.350 | 0.3% |
| Error | 316 | | | | 316 | | | | 316 | | | |

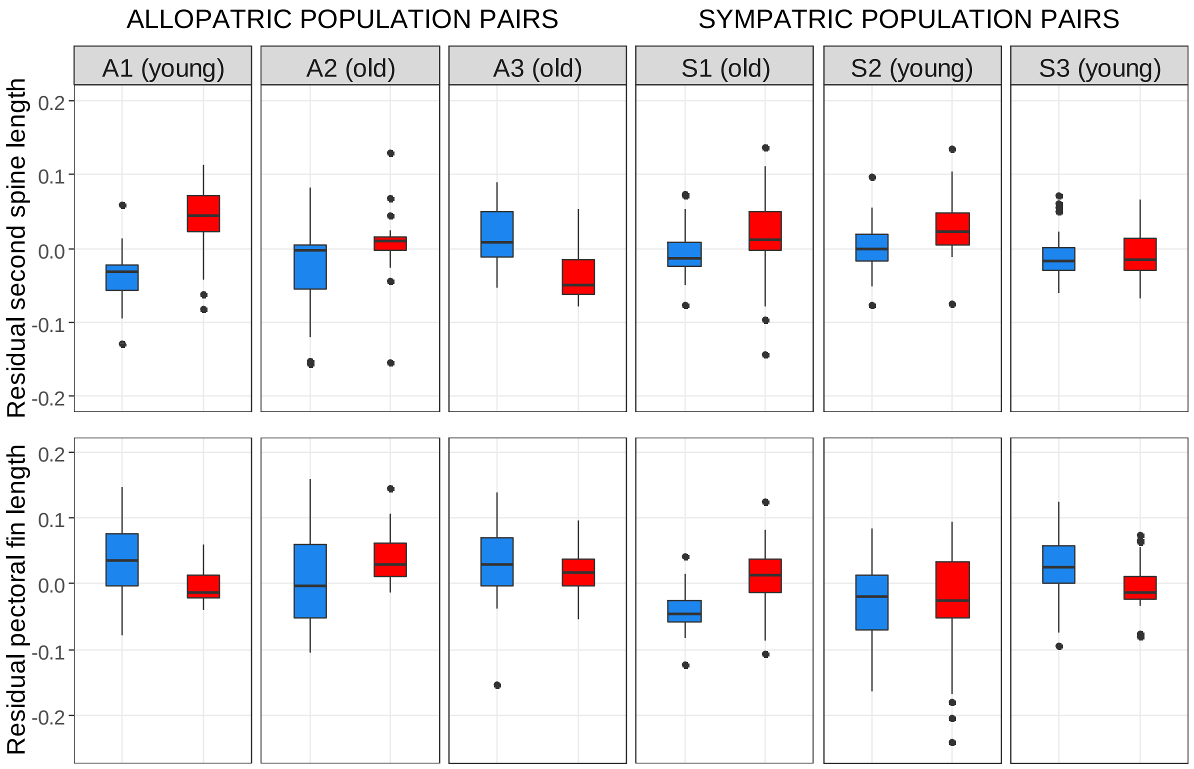


**Supplementary Figure 1.** Boxplots showing the residuals against centroid size for the second dorsal spine length and pectoral fin length. Filled circles indicate outliers. Wild-caught specimens from cold and warm habitats are indicated in blue and red, respectively. "Young" or "old" refers to the age of the warm habitat in that population pair.

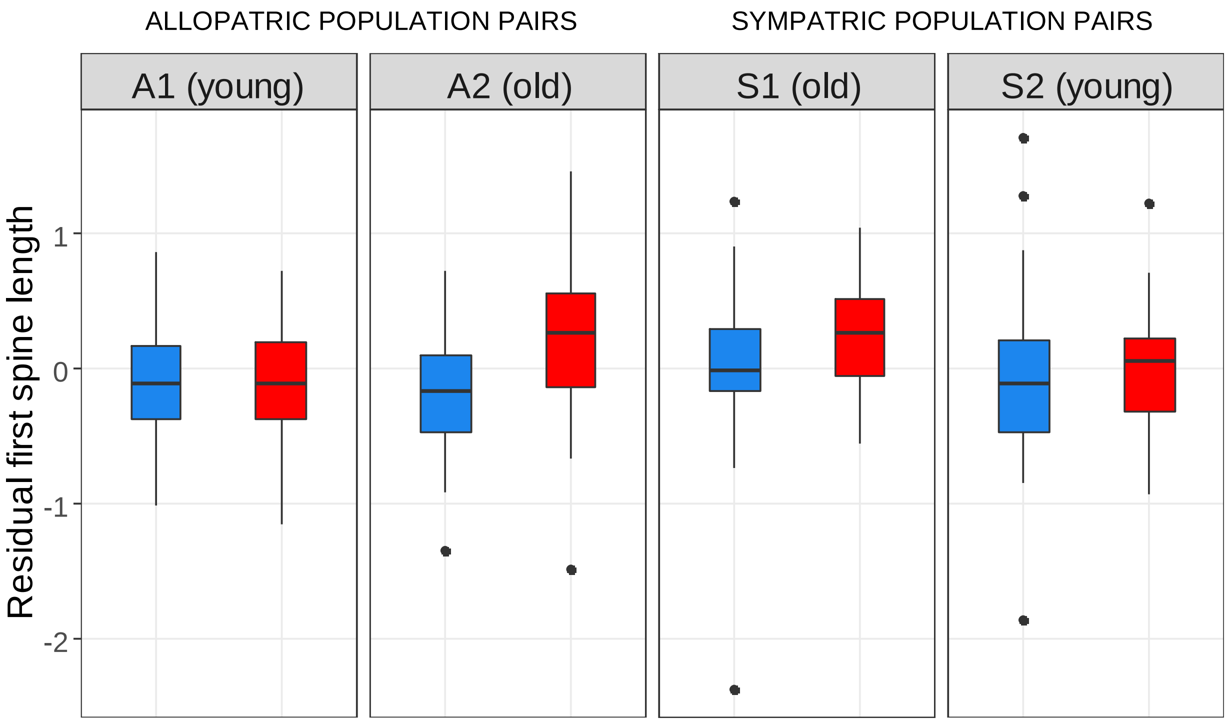


**Supplementary Figure 2.** Boxplots showing the residuals against centroid size for the first dorsal spine length in lab-reared F1 sticklebacks derived from different thermal habitats. Filled circles indicate outliers. Cold and warm thermal habitats are indicated in blue and red, respectively. “Young” or “old” refers to the age of the warm habitat in a particular population pair.

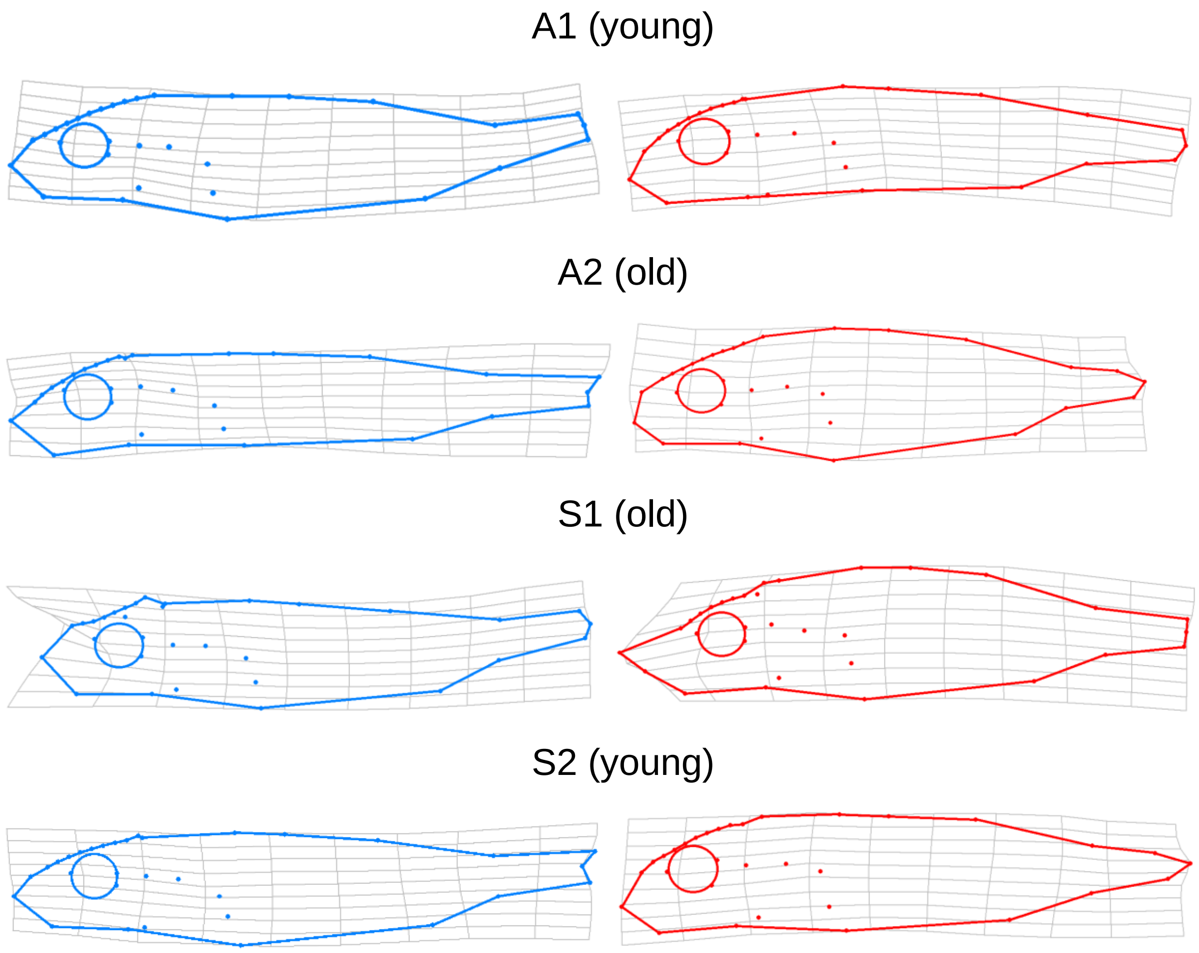


**Supplementary Figure 3.** Thin plate spline deformations showing the observed extremes in lab-reared F1 sticklebacks from four population pairs (A1, A2, S1, and S2). These fish were all reared at a common temperature (18°C). Blue and red deformations represent specimens whose parents were collected from cold and warm habitats, respectively. "Young" or "old" refers to the age of the warm habitat in that population pair. The deformation grids were generated using tpsRegr (Rohlf 2008). The shape differences were extrapolated by a factor of 3 to allow easier interpretation.
